# Oral supplementation with Nicotinamide Riboside treatment protects RGCs in DBA/2J mouse model

**DOI:** 10.1101/2024.12.03.626460

**Authors:** Nan Zhang, Ying Li, Xian Zhang, Micah A. Chrenek, Jiaxing Wang, Preston E. Girardot, Jana T. Sellers, Charles Brenner, Xiangqin Cui, Eldon E. Geisert, John Nickerson, Jeffrey H. Boatright

## Abstract

**Purpose:** The aim of this study was to test whether oral administration of nicotinamide riboside (NR), the nicotinamide adenine dinucleotide (NAD+) precursors, protect retina ganglion cells (RGCs) from neurodegeneration in DBA/2J (D2) mice, which is a widely used mouse model of age-related inherited glaucoma.

**Method:** Oral NR or NAM administration (NR low dose: 1150mg/kg; NR high dose: 4200mg/kg; NAM low dose group: 500mg/kg; NAM high dose: 2000mg/kg of body weight per day) essentially started when D2 mice were 4 or 9 months old and continued up to 12 months old. Control cohort identically received food/water without NAM or NR. Intraocular pressure (IOP) was measured every month until experiment completion. Pattern electroretinography (PERG) was recorded. Retinas were harvested for whole mount immunofluorescence staining with RGCs marker Brn3a and imaged by fluorescent confocal microscopy. Optic nerves were harvested for axon staining and quantification. Retinal NAD+ levels were enzymatically assayed.

**Results:** NR oral supplementary treatment started at 4 months old robustly increased retinal NAD+ levels in D2 mice (NR^High^ vs. vehicle: 273.7±23.59% vs. 108.70±12.10%, *p*<0.001). In aged vehicle group (12 months old), there was significantly diminution of the P1 and N2 components of PERG response compare with naïve group (naïve vs. vehicle: P1: 7.82± 0.70uV vs 1.63± 0.17uV, *p*<0.0001; N2: -13.29± 0.83uV vs. -3.22± 0.27uV, *p*<0.0001; Kruskal-Wallis test with Dunn‘s multiple comparison test). NR treatment preserved aged D2 visual function when mice were 9 and 12 months old. In addition, long-term NR high dose treatment significantly protected against total RGCs loss and optic nerve atrophy (RGC: NR^High^ vs. vehicle: 1412±62.00vs 475.2±94.68 cells/field, *p*<0.00001; axon numbers: NR^High^ vs. vehicle: 23990±1159 vs 8573±1160, n=41-53, *p*<0.0001). Furthermore, long-term NR supplementation prevent iris depigmentation and delayed IOP elevation.

**Conclusion:** NR oral supplementary treatment significantly preserved RGC and axon numbers, potentially preserves retinal function via elevated retinal NAD+ level in aged D2 mice. Interestingly, NR treatment also prevented iris atrophy, delayed IOP elevation associated with this glaucoma model. NR oral supplementation thus treated several aspects of murine pigment dispersion glaucoma. Given parallels between this model and glaucoma in human, out data indicate that NR is worth exploring as a therapeutic candidate in treatment of glaucoma.

## Introduction

As a worldwide disease leading of irreversible vision loss, glaucoma characterized by degeneration of retinal ganglion cells (RGCs) and optic nerve axons. Current evidence shows that loss of RGCs is related to the elevation of intraocular pressure (IOP), but other factors may also play a role [PMID:24825645]. However, even reduction of IOP is the only proven method to treat glaucoma, clinical studies show that glaucoma can still progress when IOP is pharmacologically managed [PMID:24825645]. There is thus an urgent need to develop IOP-independent neuroprotection treatment.

Nicotinamide adenine dinucleotide (NAD+) has been reported decline with aging in many preclinical models [PMID31412242]. NAD+ is enzymatic co-factor that is central to metabolism, found in all living cells. In metabolism, NAD+ is involved in redox reactions, carrying electrons, plays an important role in energy production and DNA repair [PMID29432159]. Studies show that NAD+ is modulated by metabolic stress in aged condition [PMID32694684]. The decline of NAD+ in aged tissue is showed associated with neuron dysfunction, indicating that it may play a direct role in maintaining neuronal health [PMID31280708]. Visional research found that RGCs mitochondrial function diminishes with age, likely due to declines in retinal NAD+ levels [PMID28858158]. Besides IOP, age is another significant risk factor of glaucoma [PMID26310165]. The dysfunction of RGCs with aging might increases susceptibility to the effects of elevated IOP [PMID28209901]. Studies have showed that increasing retinal NAD+ levels by systemic treatment with NAD+ precursor nicotinamide (NAM) protects in aged DBA/2J mouse model of inherited glaucoma [PMID 28487632; 29497468; 28858158;28209901]. Clinical trial research also reports that NAM supplementation can improve inner retinal function in glaucoma by testing photopic negative response (PhNR) parameters [PMID 32721104]. Therefore, use of NAD+ precursor has been proposed as a strategy for augment NAD+ levels, improving retinal function with aging, which potentially could be therapeutic target for glaucoma treatment by maintaining retinal NAD+ levels.

Nicotinamide riboside (NR) is presently in wide use as another NAD+ precursor, showing well tolerated and effectively boost NAD+ synthesis [PMID29599478]. It has been found that NR is highly bioavailable in mice and human to rise NAD+ levels, which been tested on various mice models and clinical trials [PMID32852543; 27721479;29211728]. NR is a naturally occurring vitamin B3, has been showed protection of damaged neurons in two potential mechanisms: preserving axonal NAD+ [PMID25908823] and boosting mitochondrial NAD+ [PMID25470550]. NAM and NR are converted to NAD+ by way of nicotinamide mononucleotide (NMN) in the salvage pathway [PMID12555668,11157928,15381699,15137942]. Simon John’s lab published data supports therapeutic benefit from systemic treatment with NAM in DBA/2J mouse model of inherited glaucoma [PMID28209901]. At dose of 2000mg/kg/day on this mice model, 93% of eyes did not develop glaucoma. It is equivalent to 9.8g/day for a 60kg human [PMID27057123]. A study about safety of high-dose NAM showed that high-dose of NAM should still be considered as a drug with toxic potential at adult doses in excess of 3 g/day [PMID11126400]. Therefore, biosafety profile should be considered when transfer NAM research achievement to pre-clinical use on human. The bioavailability and effectiveness of NR was been studied recently. Clinical trial study showed that oral NR administration was well tolerated without adverse evets, 8 days of oral NR supplementation significantly increasing NAD+ level in human blood [PMID: 29211728]. Meanwhile, as great as the potential for glaucoma treatment, it is sensible to assess multiple NAD+ precursors. We thus propose to test whether RGCs are protected in DBA/2J glaucoma mice model by systemic treatment with NR via oral administration.

## Method

### Animal

Adult DBA/2J mice from Jackson Laboratories were used in this study. Mice were housed in a 12-hour light-dark (7 AM on and 7 PM off) cycle with food and water in Atlanta VA Center for Visual and Neurocognitive Rehabilitation animal facility. All animal procedures were approved by the Institutional Animal Care and Use Committee (ACORP #: V010-18) and performed in accordance with National Institutes of Health guidelines and the ARVO Statement for the Use of Animals in Ophthalmic and Vision Research.

For short-term treated groups (Figure 1), the mice were administered NR (Niagen®, ChromaDex, Inc., Longmont, CO)/NAM (Spectrum, Cat#: 98-92-0, Gardena, CA) in water starting at 9 months old. Low dose of NR (1150mg/kg/d, NR^Low^) or High dose of NR (4200mg/kg/d, NR^High^) was dissolved in 200ml regular drinking water and changed twice per week. Similarly, (based on molar dose equivalent principle) NAM was administered in drinking water as low dose (500mg/kg/d, NAM^Low^) and high dose (2000mg/kg/d, NAM^High^) sub-groups. Vehicle group mice received same drinking water with no added NR or NAM. Mice were initially divided into treatment and vehicle group randomly. Both male and female were equally involved in this experiment.

**Figure 1.**
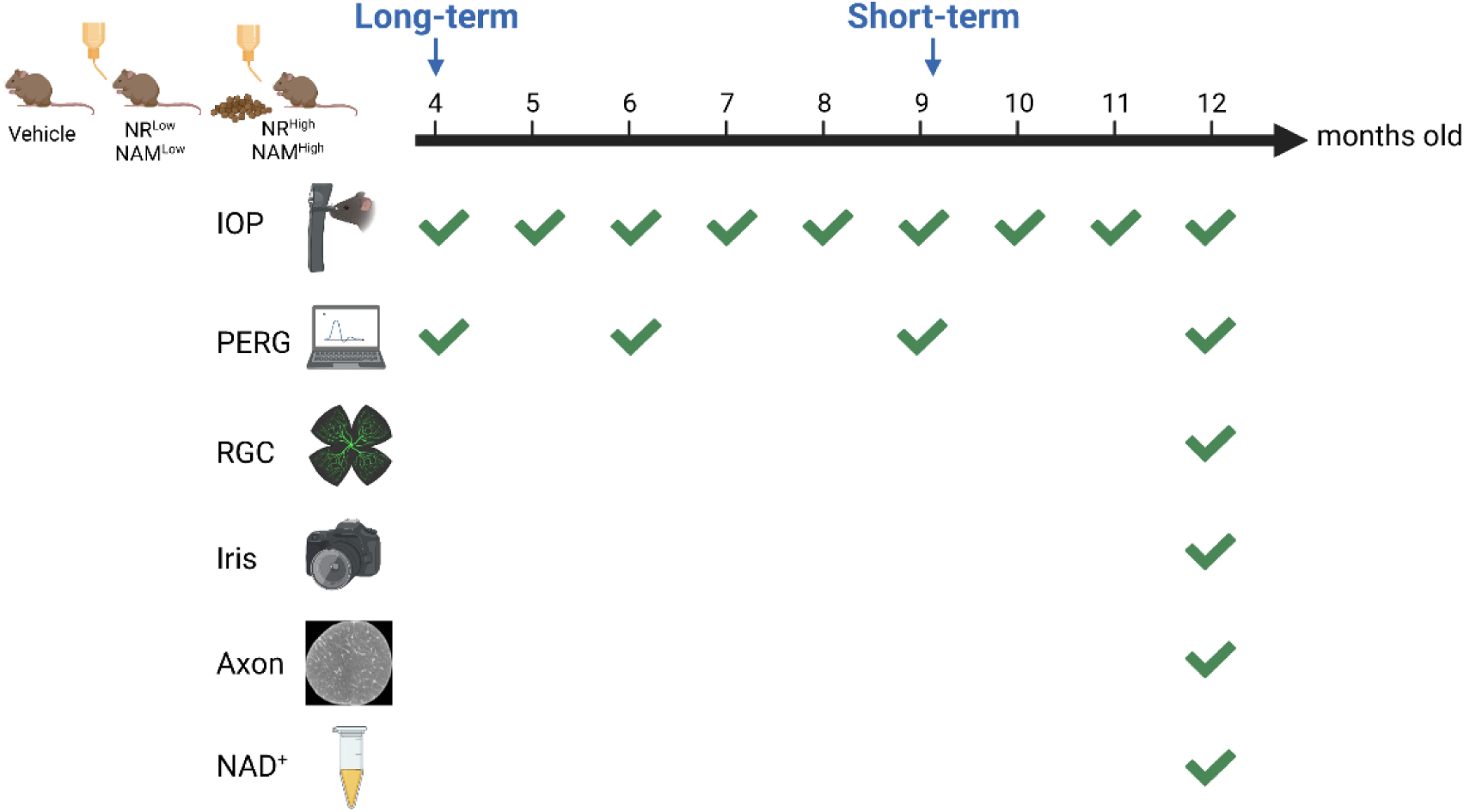
Graphic timeline. NR or NAM administration in short-term groups started when mice were 9 months old. Long-term NR or NAM treatment started when mice were 4 months old.

**Figure 2.**
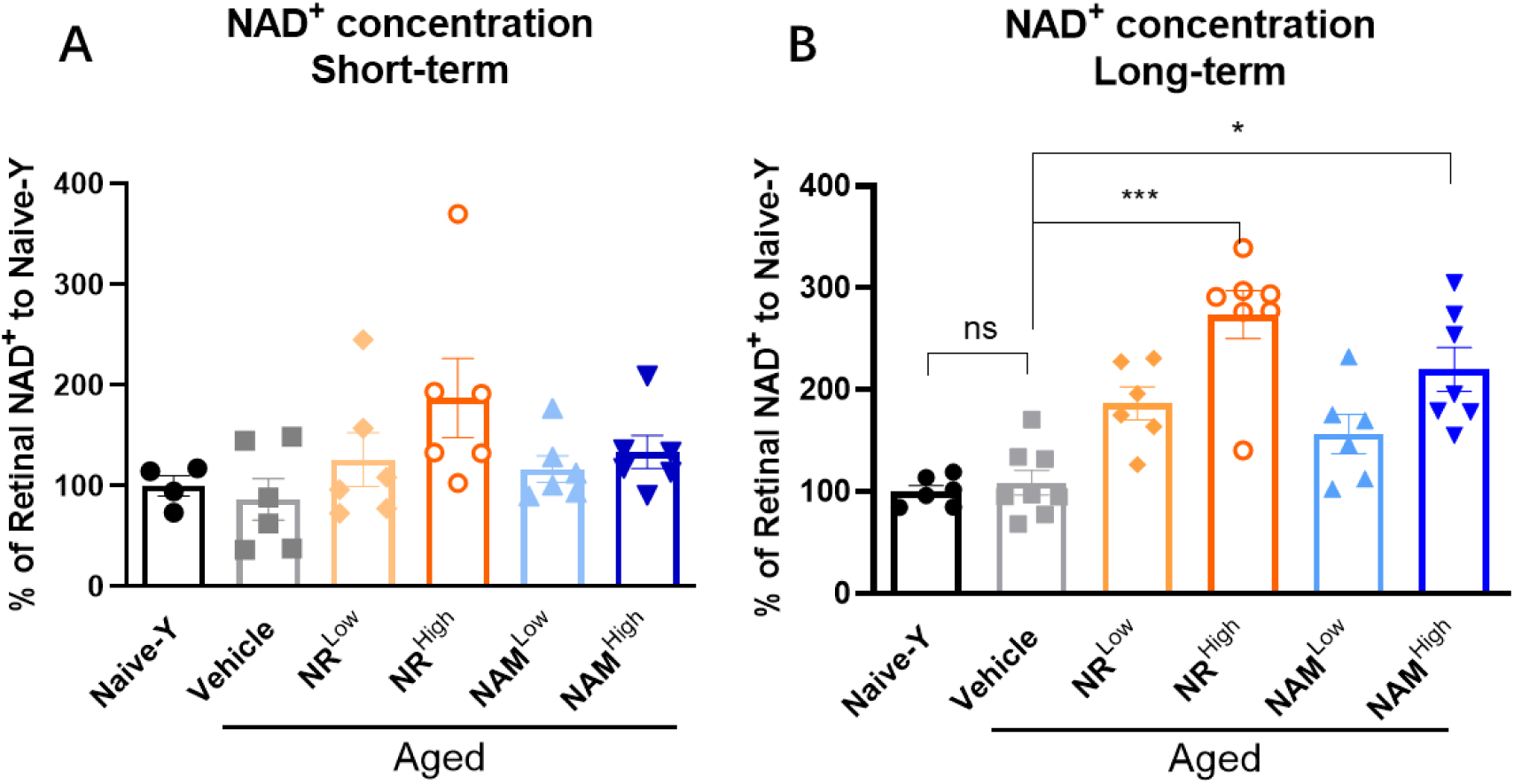
NR or NAM short-term (A) and long-term(B) oral supplementation elevated retinal NAD+ concentration. High dose of NR or NAM treatment both increase NAD+ level in retina of aged D2 mice significantly. Naïve-Y: 4 months old DBA/2J mice without any treatment. The results are represented as mean ± SEM. Kruskal-Wallis test with Dunn’s multiple comparison test, ***p <0.001, * p<0.05.

For long-term treated groups (Figure 1), the mice in low dose treated groups were administered NR (Niagen®, ChromaDex, Inc., Longmont, CO)/NAM (Spectrum, Cat#: 98-92-0, Gardena, CA) in water starting at 4 months old, which is same as short-term groups. Mice in high dose treated groups were treated with NR or NAM in both food (NR: 5G35, TestDiet; NAM: 5G34, TestDiet) and drinking water. Vehicle group mice received same food (5W80, TestDiet) and drinking water with no added NR or NAM. Mice were initially divided into treatment and vehicle group randomly. Both male and female were equally involved in this experiment.

### IOP Measurement

IOP was measured with TonoLab tonometer which is widely used for mouse in lab (PMID29421330). A rebound tonometer (Tonolab Colonial Medical Supply, Londonderry, NH) was used to measure the IOP under anesthesia with 5% Isoflurane (NDC 66794-017-25; Bethlehem, PA, USA). Same researcher in lab measured IOP every month. Measured values are averaged values of 3 repeated measurements per animal and time point. IOP readings obtained with the Tonolab instrument have been shown to be accurate and reproducible in various mouse strains, including DBA/2J (Nagaraju et al., 2007; Saleh et al., 2007).

### Pattern-ERG

Mice were dark-adapted overnight before pattern electroretinograms (PERGs) were performed. In preparation for the PERGs, mice were anesthetized using intraperitoneal (IP) injections of ketamine (80 mg/kg) and xylazine (16 mg/kg). Once anesthetized, proparacaine (0.5%; Akorn Inc.) and tropicamide (0.5%; Akorn Inc.) eye drops were administered to reduce eye sensitivity and dilate the pupils. Mice were placed on a heating pad (37 °C) under dim red light provided by the overhead lamp of the Celeris-Diagnosys system (Diagnosys, LLC, Lowell, MA, USA). Transient PERG responses were recorded using black and white vertical stimuli delivered on a Celeris system. The pattern stimulator was placed in contact with tested eye. The flash stimulator for contralateral eye acts the reference electrode. Briefly, pattern stimuli of 50 cd·s/m^2^ were presented and 600 averaged signals with cut-off filter frequencies of 1 to 300 Hz were recorded under scotopic conditions. P1 value is measured from the N1 trough to the peak of P1, N2 is measured from the preceding P1 peak to the nadri of N2 (as shown in figure 2A). Each mouse was placed in its home cage on top of a heating pad (39 °C) to recover from anesthesia.

### NAD+ Concentration Measurement

NAD+ Concentration Measurement methods had been reported by our group before [PMID: 32852543]. Levels of NAD^+^ in homogenates were measured following manufacturer’s instructions (Abcam; ab. 65348; #Lot: GR3226737-3; San Francisco, CA). In brief, retina samples were homogenized in extraction buffer. Extracted samples were separated into two aliquots. One was used to measure total NAD (NADt). For NADH specific measurements, samples were heated to 60°C for 30 min to decompose NAD^+^. Extracted samples were placed in a 96-well plate and the NADH developer was added into each well. The plate was placed into a hybrid reader and read every 30 min at OD 450 nm while the color was developing. Data from the 2-hour timepoint are presented. NADt and NADH concentrations were quantified against an NADH standard curve. NAD^+^ was calculated with the equation NAD^+^ = NADt - NADH.

### Histology: Retina whole mount

For retina flat mount preparation, mice were deeply anesthetized with ketamine (80 mg/kg) and xylazine (16 mg/kg), then perfused through the heart with 0.9% saline (Thermo Fisher Scientific; Lot# 130586; Waltham, MA, USA) followed by 4% paraformaldehyde (Electron Microscopy Sciences; Lot# 191203-27; Hatfield, PA, USA). The dissected eyeballs were post-fixed in 4% paraformaldehyde for an extra hour at room temperature. Retinas were removed from the globe and cut into quarters, creating four flaps or petals. Then, retinas were rinsed in PBS with 1% Triton X-100(Sigma-Aldrich, Cat. # 9002-93-1, St. Louis, MO, USA) and blocked with 5% normal donkey serum in a 96-well plate, and placed in primary antibody, Brn3a+ (1:1000; Santa Cruz; SC-31984; Dallas, TX, USA) overnight at 4 °C. The whole mount retinas were washed in PBST (PBS with 0.1%Tween-20) for 3 times and then placed in secondary antibody (1:1000, Alexa Fluor 488 AffiniPure Donkey Anti-Goat, Invitrogen, Cat# ABS82, Eugene, OR, USA). After 3 washes in PBST, cover slips were placed over the retina whole mounts using Fluoromount - G (Southern Biotech, Cat. # 0100-01, Birmingham, AL, USA).

Retinal flatmounts were imaged with a Nikon Eclipse Ti confocal microscope (Nikon, Inc., Melville, NY, United States) under 20x magnification. Images were processed with a custom pipeline in CellProfiler (https://cellprofiler.org/citations) to count the number of Brn3a+ labeled cells in a masked manner [PMID27119563]. For statistical analysis, the immune-positive RGCs per retina was compared.

### Analysis of optic nerve axon damage

Mice were anesthetized with ketamine (80 mg/kg) and xylazine (16 mg/kg) and perfused through the heart with saline followed by 2% paraformaldehyde and 2% glutaraldehyde in phosphate buffer (pH 7.4). The optic nerves were carefully dissected and post-fixed for 24 hours at 4 °C. Nerves were dehydrated and embedded in plastic (Libby et al., 2005). Thin sections (0.7 *μm*) were cut from each nerve and mounted on glass slides. Sections were stained using a modified Toludine Blue staining protocol. Slides of optic nerve axon sections were imaged with a Nikon Eclipse Ti confocal microscope (Nikon, Inc., Melville, NY, United States) under 60x magnification.

The number of axons per nerve from all groups of mice were automatically calculated using AxoNet software (PMID32415269). Optic nerves were analyzed and determined to 3 different damage levels: No or early damage (NOE) – less than 20% axons damaged; Moderate damage (MOD) – more of 20% axon loss and less than 50%; Severe (SEV) – more than 50% axonal loss and damage.

### Anterior chamber observation and Iris atrophy grading

Clinical presentation of depigmenting iris disease of 1-year old DBA2J mice were observed under slit lamp. Iris transillumination photographic images were taken before experiment completion. Pathological changes in Iris were assessed based on transillumination assay, and grading into 4 different sub-groups with order from Level 0: no iris atrophy; Level 1: mild iris atrophy; Level 2: moderate iris atrophy; to Level 4: severe iris atrophy. Every level of iris depigmenting group from various treatment group was counted as showed in table chart (Figure 6).

### Statistical analyses

The personnel conducting assessments in experiments that required judgement were masked to the specific treatment group from which sampling arose. This included semi-automated marking of PERG peaks and nadirs, semi-automated counting of axon numbers in optic nerve sections. Statistical analyses were conducted using Prism 8.4.2 Software (GraphPad Software Inc. La Jolla, CA, USA). Kruskal-Wallis test with Dunn‘s multiple comparison test was performed for biochemical and morphometric data. Chi-square statistic were used for discontinued data type. Unless otherwise noted, n is the number of animals per experimental group. For all analyses, results were considered statistically significant when *p*< 0.05. Data are presented as mean ± SEM.

## Results

### 1. Retinal NAD+ levels in aged DBA/2J increases with NR oral supplementation

To determine whether oral supplementation of NR could elevate NAD+ level in mouse retina, we harvested and assayed mice retina tissue from different treated groups for NAD+/NADH content. As shown in figure2, compare with aged vehicle group (12 months old), both low and high doses of NR and NAM with long/short-term treatment elevated NAD+ level in various degrees. Especially in long-term treated groups, NAD+ level from both NR and NAM high dose groups showed statistically significant elevation compared to the vehicle group (NR^High^ vs. vehicle: 273.7±23.59% vs. 108.70±12.10%, *p*<0.001; NAM^High^ vs. vehicle: 220.10±21.54% vs. 108.70±12.10%, *p*<0.05; n=6-7, Kruskal-Wallis test with Dunn‘s multiple comparison test). However, low dose groups did not show statistically significant NAD+ level elevation compared with vehicle group.

### 2. NR partially preserved visual function assessed by PERG in aged DBA/2J mice

PERG data were collected to assess RGC functional changes in aged D2 mice. Figure 3A showed the representative waveforms of PERG from vehicle group and NR or NAM treated groups. Quantification of PERG amplitudes from short-term groups was presented in figure 3B.

**Figure 3.**
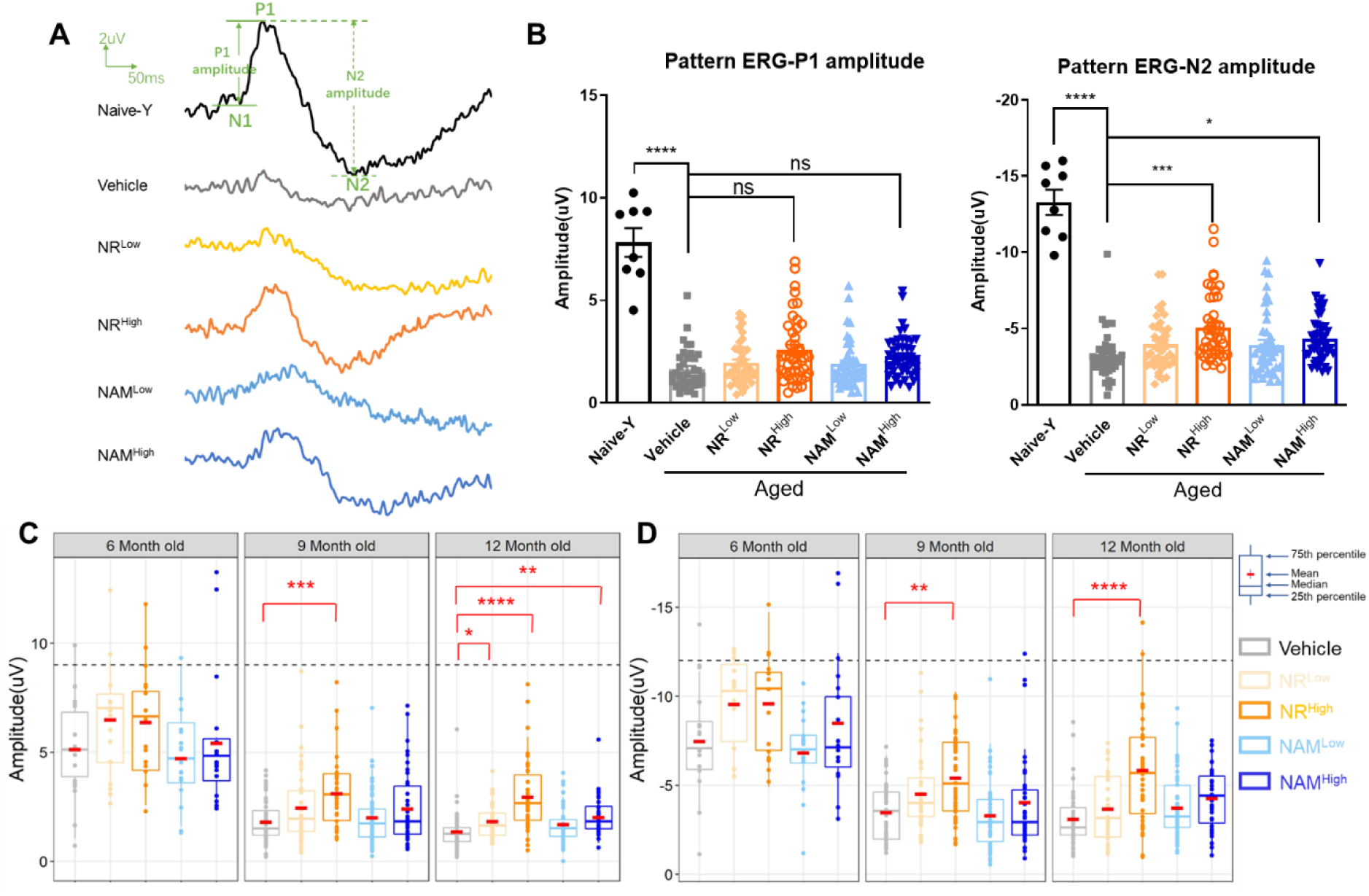
NR or NAM oral supplementation preserve RGC function in aged D2 mice. A. Representative pattern ERG wave-forms from each group and measurements of P1 and N2 amplitudes. B. Quantification of P1 and N2 amplitudes from short-term NR/NAM treated groups. C-D. Quantification of P1 and N2 amplitudes from long-term NR/NAM treated groups. Black dash lines in C&D represent the mean amplitude of Naïve-Y group at 4 months old. The results in figures are represented as mean ± SEM. Kruskal-Wallis test with Dunn‘s multiple comparison tests were conducted between the mean amplitudes in all combinations. * p<0.05, ** p<0.01, *** p<0.001, **** p<0.0001. n=8-44.

Compare with young naïve D2 mice (4 months old), aged vehicle ones had obvious decline on both N2 and P1 amplitudes (P1: 7.82± 0.70uV vs 1.63± 0.17uV, *p*<0.0001; N2: -13.29± 0.83uV vs. -3.22± 0.27uV, *p*<0.0001; n=8 in naïve group, n=36 in vehicle group, Kruskal-Wallis test with Dunn‘s multiple comparison test), indicating glaucomatous function loss appeared in aged D2 mice.

Short-term oral NR or NAM supplementation could attenuate the functional loss in aged D2 mice. N2 amplitudes from NR or NAM high dose group both showed significant improvement compare with aged vehicle group (NR^High^ vs. vehicle: -5.05±0.33uV vs. -3.22±0.27 uV, *p*<0.001; NAM^High^ vs. vehicle: -4.33±0.24 vs. -3.22±0.27 uV, *p*<0.05; n=36-44, Kruskal-Wallis test with Dunn‘s multiple comparison test). However, no significant difference been found between NR^Low^/ NAM^Low^ and vehicle group.

For long-term groups, PERG was performed when mice were 4, 6, 9 and 12 months old (Figure 3C&D). For mice in vehicle group, both P1 and N2 amplitudes diminished slightly at 6 months old, and significantly decreased at 9 months old compare with naïve group (P1: Naïve-Y vs. vehicle-9m: 8.63±0.58uV vs. 1.80±0.15uV, *p*<0.001; N2: Naïve-Y vs. vehicle-9m: - 11.66±0.57uV vs. -3.49±0.25uV, *p*<0.001). The visual function was further damaged when vehicle mice were 12 months old (P1: Naïve-Y vs. vehicle-12m: 8.63±0.58uV vs. 1.37±0.13uV, *p*<0.001; N2: Naïve-Y vs. vehicle-12m: -11.66±0.57uV vs. -3.11±0.24uV, *p*<0.001).

Compare with vehicle group, NR high dose treatment preserved visual function significantly (9 months old: NR^High^ vs. vehicle:3.11±0.26uV vs. 1.80±0.15uV, *p*<0.001; NR^High^ vs. vehicle: -5.42±0.37uV vs. -3.49±0.25 uV, *p*<0.001;12 months old: NR^High^ vs. vehicle: 2.90±0.27uV vs. 1.37±0.13uV, *p*<0.001; NR^High^ vs. vehicle: -5.85±0.50uV vs. -3.12±0.24 uV, *p*<0.001). NAM high dose treatment also partially protected visual function loss when mice were 9 months old, but failed to show protection at 12 months old (9 months old: NAM^High^ vs. vehicle: 2.38±0.25uV vs. 1.80±0.15uV, *p*<0.001).

### 3. NR treatment aids more RGC survival in aged DBA/2J

RGC survival was assessed by counting Brn3a labeled cells. For short-term groups, as shown in figure 4B, aged D2 mice suffered significant RGC loss compared with young D2 mice (Naïve-Y vs. vehicle: 1700 ±14.44 vs 503.40±146.20 cells/retina, n=11-12, *p*<0.0001). However, high dose NR or NAM treatments partially but not significantly protected against total RGCs loss in retinal immunofluorescent Brn3a+ cell counts. Both NR and NAM low dose groups did not show protection compare with vehicle on Brn3a+ RGCs.

**Figure 4.**
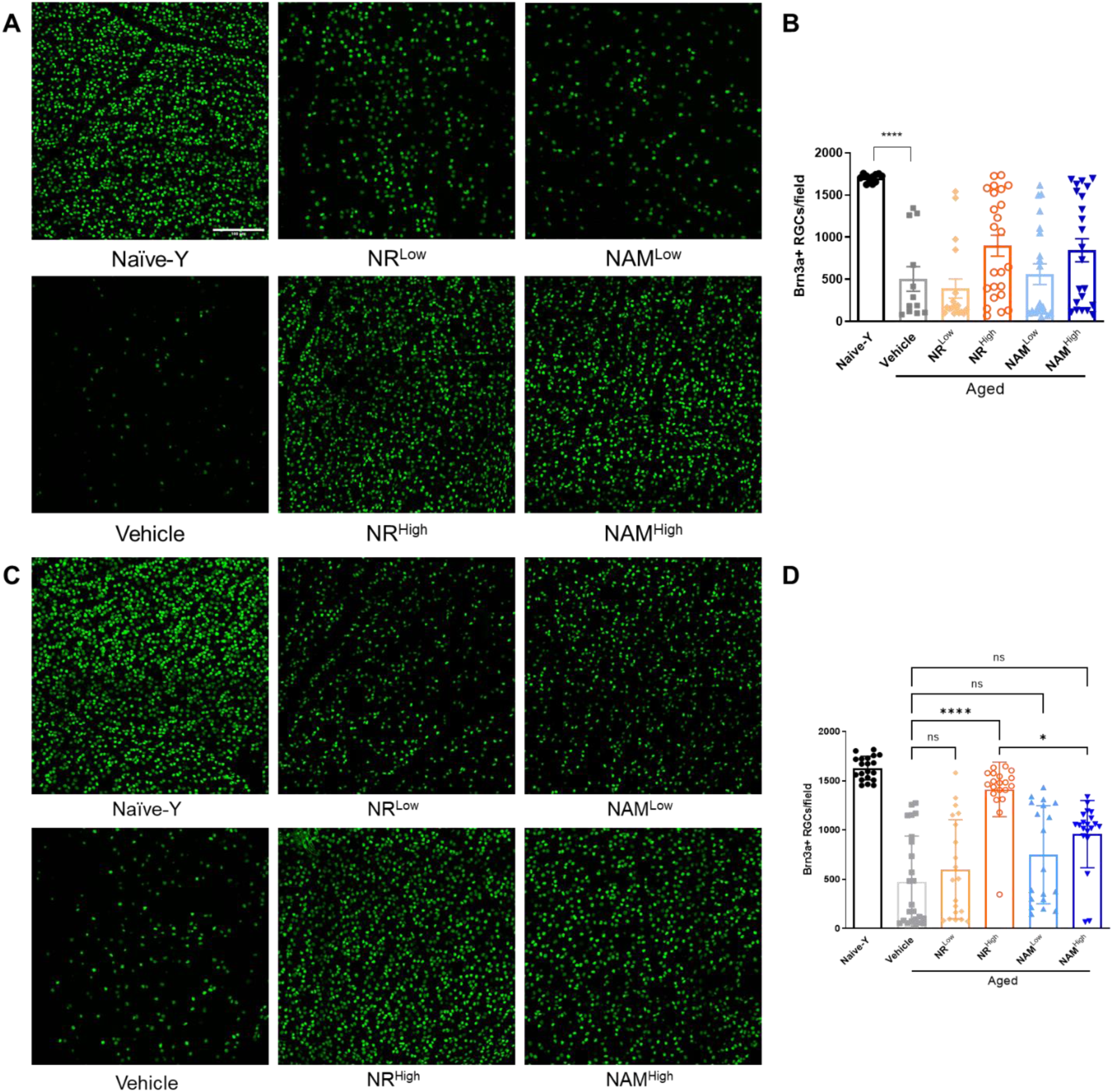
NR oral supplementation partially protected against total RGCs loss in retinal immunofluorescent Brn3a+ cell counts. A, C. Representative images of retina flatmounts stained against Brn3a+ from different short-term(A) and long-term(C) treated groups. Images were chosen randomly from the region which distance to the optic nerve was 1.0 mm. B, D. Quantification of Brn3a-positive cells, assumed to be RGCs for short-term (B) and long-term groups(D). The results are represented as mean ± SEM. Kruskal-Wallis test with Dunn‘s multiple comparison test, * p<0.05, **** p<0.0001. n>12 in each group. Scale bar: 100 µm.

Long-term NR high dose treatment significantly aid more RGC survival in aged D2 mice (Figure 4D, NR^High^ vs. vehicle: 1412±62.00vs 475.2±94.68 cells/field, n=20-24, *p*<0.00001). NAM high dose group also showed trend of attenuation on RGC loss compare with aged vehicle group, but the difference was not statistically significant (NAM^High^ vs. vehicle: 958.8±76.38 vs 475.2±94.68 cells/field, n=22-24, *p>0.05*). Notably, NR^High^ group showed more effctive protection on RGC survival compare with NAM^High^ (NR^High^ vs. NAM^High^: 1412±62.00 vs 958.8±76.38 cells/retina, n=20, *p*<0.05).

### 4. NR treatment significantly protected optic nerve axons from loss in aged DBA/2J mice

Axon numbers were quantified by using AxoNet in optic nerve sections stained with toluidine blue. For short-term groups, as shown in figure 5C, there is a significant decrease in axon number in vehicle group compare with Naïve-Y group (Naïve-Y vs. vehicle: 18100±1283 vs. 5820±1036, p<0.0001). Treatment with NR and NAM high dose partially prevented the axon loss in glaucomatous eyes but the difference was not significant. To further analyze the protective effects of NAM or NR on axon survival, we divided optic nerve sections into three damage levels: none or early (NOE), moderate (MOD) and severe (SEV). The representative images from different level were shown in figure 5A. NR high dose treatment reduced the percentage of severe damaged nerves and increased the percentage of moderate damaged optic nerves compare with vehicle group (Figure 5B). However, the interventional treatment with NR or NAM at 9 months old (when the D2 eyes already have had IOP elevation) did not show significant protection from optic nerve degeneration as assessed by axon stain.

**Figure 5.**
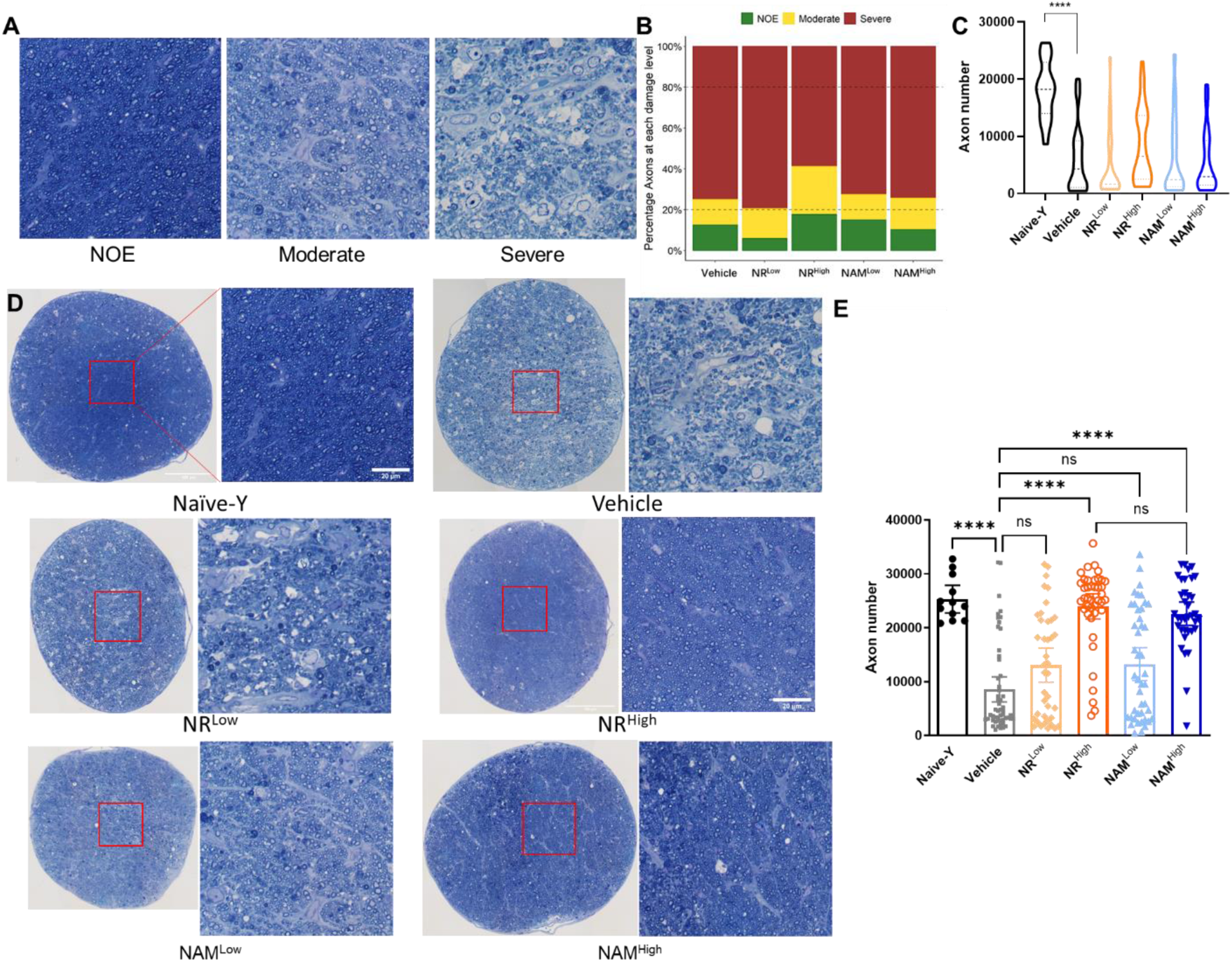
NR or NAM oral supplementation prevent axon loss in aged D2 mouse. A. Representative axon images from each damage level with toluidine-blue staining. B. Percentage axons at each damage level. Green: none or early damage [=<20% axon loss; none or early (NOE)]; yellow: moderate damage (20-50%axon loss); red: severe damage (=>50% axon loss). C. Quantification of survival axon number using AxoNet for short-term groups. D. Representative axon images from each long-term treated groups. E. Quantification of axon number for long-term groups. Kruskal-Wallis test with Dunn’s multiple comparison test (C&E), ****p<0.0001. Chi-square statistic(B). n>31 in each group.

For long-term groups (Figure 5D, E), treatment with high dose NR or NAM did significantly prevent the axon loss in aged D2 mice (NR^High^ vs. vehicle: 23990±1159 vs 8573±1160, n=41-53, *p*<0.0001; NAM^High^ vs. vehicle: 22627±1076 vs 8573±1160, n=36-53, *p*<0.0001). NR and NAM low dose treatments didn’t show statistical protection on axon numbers.

### 5. NR delayed IOP elevation and protected iris from depigmentation in aged DBA/2J

IOP elevation was observed in long-term NR and NAM treated groups per months. Compared with IOP of Naïve D2 mice (4 months old, black dash line in figure 6A), mean IOP of vehicle mice started to elevate at their 7 months old, and reached the peak at 9 months old. Moreover, NR intervention affect the IOP elevation in aged D2 mice, that there is significantly different in IOP assessment between NR high dose treated and vehicle eyes at the timepoints of 7 and 9 months old (*Kruskal-Wallis test with Dunn’s multiple comparison test*, p<0.01). NAM treated groups also had mild trends of delayed IOP elevation, but there were no statistical differences between NAM treated groups and vehicle at each timepoints.

**Figure 6.**
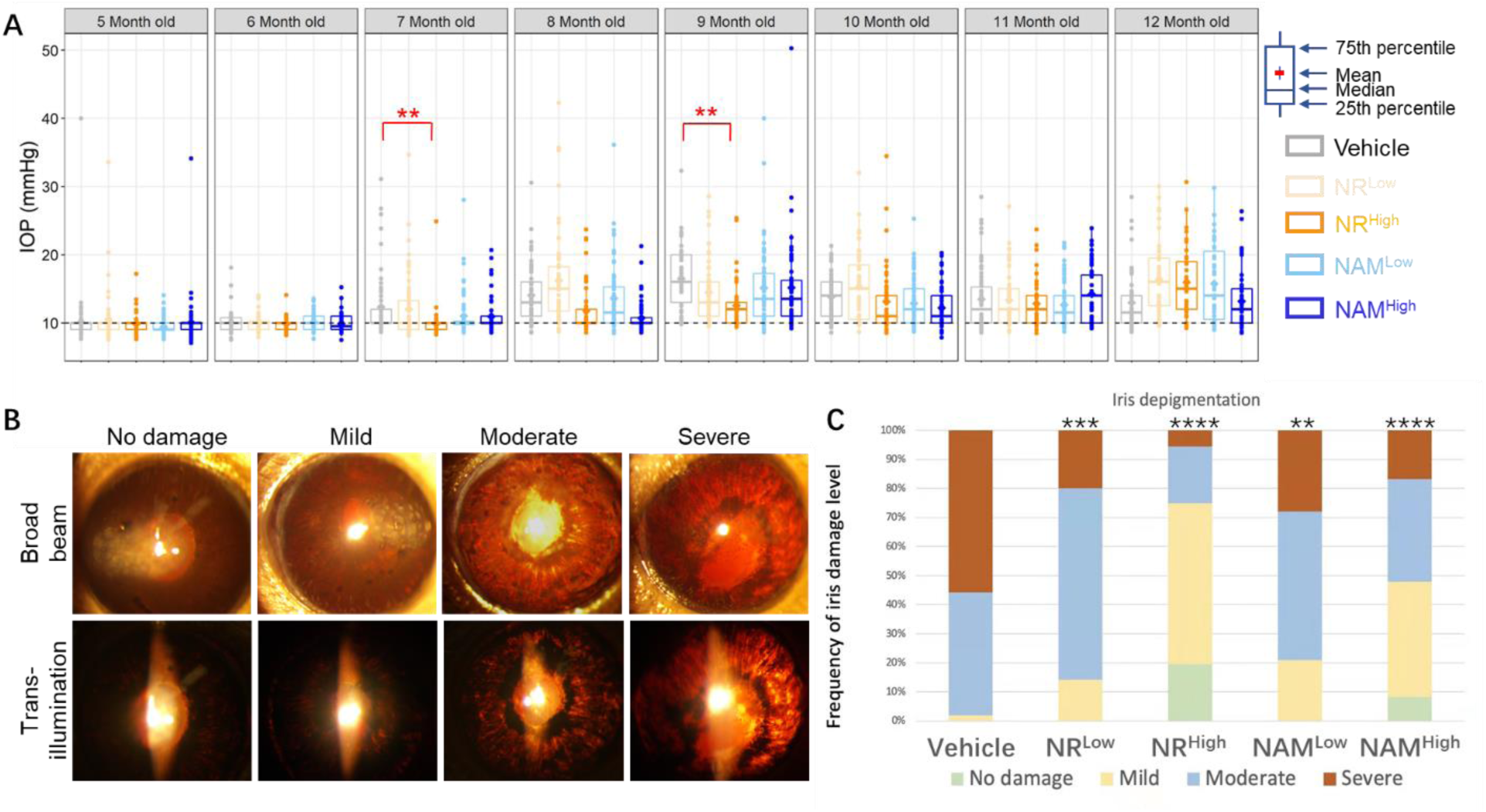
NR oral supplementation delayed IOP elevation and protected iris from depigmentation in D2 mice. A. IOP from each long-term treated group for D2 mice. Black dash line: mean IOP value from Naïve-Y group. B. Representative images for each damage level of iris for aged D2 mice. C. Percentage iris at each damage level for each group. Kruskal-Wallis test with Dunn’s multiple comparison test (A), **p<0.01. Fisher’s exact test (C), comparisons were done between vehicle and each other group, **p<0.01, ****p<0.0001. n>38 in each group.

To further investigate the potential reasons why NR treatment delays IOP elevation in D2 mice, we observed the iris pathological changes for each mouse when they were 12 months old. According to the observation of anterior chamber and trans-illumination representation, the levels of iris atrophy were graded into four levels, no damage, mild, moderate and severe. The representative images from different level were shown in figure 6B. *Fisher’s exact statistic* test was performed to analyze the effects of NR or NAM on iris depigmentation in aged D2 mice (Figure 6C). Results showed that NR or NAM treatments protected iris from depigmentation respectively in various degree, which might explain the delayed IOP elevation for D2 mice.

## Discussion

Recently supplementation with NR or NAM, as major NAD+ precursors, has been shown neuroprotection benefits on eye health, and neurodegenerative disorders. [ PMID: 31185090, 29497468, 26118927] In present study, we orally treated aging DBA/2j mice which mimic human hereditary glaucoma, with NR or NAM-rich food/water and tracked the functional and morphological changes. Our work results showed that long term oral supplementation with NR preserved RGC function as well as promoted RGC and axon survival, caused by glaucomatous damage in D2 mice. NR also protected iris from depigmentation and delayed IOP elevation during aging of D2 mice. NR interestingly showed more efficacy on neuroprotection on D2 glaucoma mouse model comparing with NAM at equivalent molar dose, although both the neuroprotective effect of NR and NAM were dose dependent.

Evidence has been shown that NAD+ is critical in retinal metabolism. NAD+ deficiency has been found related to retinal degenerative disease in multiple human diseases and mouse models [PMID: 22842229, 29184169, 29674119, 27681422]. The role of NAD+ in glaucoma also has been discussed by many research groups. Both clinical trials and basic science research on rodent model reported that decline of NAD+ contributed to the susceptible to glaucomatous neurodegeneration [PMID: 31185090, 27721479]. Lower NAM level in POAG patients and NAD+ level in aged D2 mice retinal tissue both indicated that insufficiency of NAD+ level might be contributed to susceptibility to glaucoma [PMID: 31185090]. Thus, NAD+ precursors are considered as promising therapeutic candidates for glaucoma [PMID: 32961812, 28858158]. As two important NAD+ precursors, NR and NAM convert to NMN through salvage pathway via NRK1 or NRK2 and Nicotinamide phosphoribosyltransferase (Nampt), respectively to increase NAD+ synthesis [PMID: 15137942]. We thus examed the NAD+ levels in aged D2 mouse retina using NAD+ concentration assays and found that long term orally supplementation with NAD+ precursors significantly increased retinal NAD+ levels. NAD+ elevation effect from NR and NAM was related to the daily intake dose, and the effect of increasing retinal NAD+ from NR oral supplementation showed higher efficacy than equivalent molar dose of NAM oral supplementation, which is similar as research from Brenner and Canto’s group, in which they used intraperitoneal injection. [PMID: 27725675]. Samuel et.al compared the orally bioavailability of NR and NAM in mice and human. In their study, they calculated the total NAD+ under curve area in liver in 12 hours after oral administration of NR or NAM, the results indicating that NR raised NAD level more rapidly and effectively than NAM [PMID: 27721479].

Theoretically NR shows more paths to generate NAD+, that besides NRKs pathway, NR could also be converted to NAM by purine nucleoside phosphorylase, then produce NMN, finally generate NAD+ [PMID: 31795381].Our data from late start treatment cohorts also showed obvious trend of increasing retinal NAD+ content after short term of NAD+ precursors oral supplementation, especially NR high dose group, however, the elevation compared with vehicle cohort did not reach statistically significant level (Figure 2B). It might be due to relative high variation of concentration assays we used, which remind us of should increase sample size when checking NAD+ concentration assay.

Aging DBA/2j mice develop progressive ocular abnormalities including iris pigment dispersion, iris atrophy, anterior synechia and hyphema, along with gradual elevation of IOP. Significant iris pigment epithelium loss and widespread transillumination of the iris usually occur when they are 6-7 months old. [PMID9579474] In our study we observed the anterior chamber of NR or NAM treated mice and found the level of iris transillumination and incidence of hyphema were significantly lower than vehicle cohort. Our data indicated that NR also protect iris from atrophy pathological degeneration, delayed the IOP elevation in D2 mice in our study (Figure 6). Currently the mechanism that why NR-treated D2 showed resistance to typical iris pathological changes in aged D2 mice is not convincingly elucidated in this study. However, there are some possibilities might be investigated in future. Firstly, NAD is central to energy metabolism, serving as an essential coenzyme in glycolysis [28792876]. Iris pigment dispersion occurs in D2 mice due to a mutation in the glycoprotein (transmembrane) Nmb gene (Gpnmb^R150X^), but was not observed in the control strain D2-Gpnmb+/SjJ mice [35185449]. Furthermore, Shimazaki’s group reveal that down-regulation of glycolysis, NAD/NADH metabolism lead to mitochondrial degeneration. [32426498] These defect possibly were corrected to increase NAD+ levels and improve mitochondrial function by NR or other NAD precursors as many studies showed [22682224] Secondly, it has been discussed that iris pigment epithelial cells are similar as retinal pigment epithelium (RPE). They have the same embryonic origin --neural crest cells of embryological optic cup. Thumann demonstrated that in vitro cultured IPE cells do acquire functions, such as specific phagocytosis of rod out segments, that are characteristic of RPE cells, and have been shown to have the potential to carry out many functions like retinal metabolism [11166346]. Study indicated that in RPE cells derived from human pluripotent stem cells, the TGF-beta pathway was inhibited after treatment with NAM.[PMID:28132833] Therefore, NR might play a role to reduce the inflammatory response in anterior chamber and iris tissue of aged D2 mice. Similar as Arnhold pointed out [PMID15144867], we assume that iris pigment epithelium cell is a possible cell source for future treatment of neurodegenerative diseases as like glaucoma. Our data strongly support that NR could be promising protector for iris pigment epithelium cells in iris degenerative diseases.

The most encouraging part of our data is that long term of NR oral supplementation robustly prevents all assessed signs of glaucoma including RGCs loss and optic nerve axon degeneration in D2 mice (Figure 4&5). Simon John’s group reported that oral administration of NAM protected visual function loss, RGC death and axon degeneration from glaucomatous damage in D2 mice [PMID: 31185090, 29497468]. NAM-rich diet preserved RGC function by increase amplitudes of flicker-PERG as well as increased mitochondrial intensity in aged D2 mice [PMID: 32605122]. Clinical trial demonstrated that NAM was well tolerated to human and 3 months of NAM supplementation could significantly improve inner retinal function in glaucoma patients [PMID: 32721104]. Study from Takagi’s group suggested that intravitreal injection of NR protected against axonal loss in tumor necrosis factor-induced optic nerve degeneration model via upregulation of nicotinamide riboside kinase 1 (NRK1) and SIRT1-autophagy pathway [PMID:32820458]. Previously our group also published that systemic NR treatment by i.p. injection was neuroprotective in both optic nerve crush and microbead injection (ocular hypertension) mouse model [REF:34208613]. However, direct comparison between NR and NAM using glaucoma mouse model has not been reported yet. In present study both NR and NAM were set equivalent molar dose into separate treatment groups and the data supported that NR was more efficient and more effective compared to NAM showing the RGC and axon neuroprotection benefit in aged D2 mice. NRKs are rate limiting kinase for NR converting to NMN to generate NAD+, while NRK genes were lowly expressed in RGC as shown in RNA-seq datasets published previously [PMID: 31185090]. However, we found NRK1 expressed in RGCs via immunostaining [supplementary figure?], as same as Takagi’s group [PMID: 32721104].

It is not clear of mechanism that how NR promote survival of RGCs and axons in glaucoma, but makes exogenous NR administration possible, reasonable by boosting NRK1 levels and accelerating NAD+ biosynthesis in RGCs. The mechanism of this regulation can be further investigated via up-regulation or down-regulation of key enzyme, deserve to be explored in future.

## Acknowledgements

We thank ChromaDex, Inc. for supplying the NR.

## Support funding

VA I01RX002806 (JHB); VA I21RX001924 (JHB); VARR&D C9246C (Atlanta VAMC); Abraham J and Phyllis Katz Foundation (JHB); NIH R01EY028859 (JHB); NIH P30EY06360 (Emory); Owens Family Foundation (EEG); Research to Prevent Blindness (Emory), BrightFocus Foundation G2020286 (JHB&YL).

## Notes

### Competing Interest Statement

The authors have declared no competing interest.

